# Microbiome enrichment from a wild relative improves Black Soldier Fly larval survival, pathogen suppression and growth

**DOI:** 10.1101/2025.10.27.684714

**Authors:** Hauke Koch, Amy M Paten, Matthew J Morgan

## Abstract

Captive rearing can disrupt animal microbiomes, reducing diversity and impairing host performance. For the larvae of Black Soldier Flies *(Hermetia illucens*; BSFL), a key species in waste bioconversion, microbiome engineering offers a strategy to enhance growth and disease resistance. We tested whether enriching the BSFL microbiome with microbes from a wild relative, *Exaireta spinigera*, improves larval performance. Axenic BSFL neonates were inoculated with (1) a captive BSFL microbiome, (2) an *E. spinigera* microbiome, (3) a mixed inoculum of the two, (4) an environmental, diet derived microbiome, or (5) remained axenic. After two weeks, larvae receiving *E. spinigera* microbes (alone or mixed) exhibited significantly higher growth and survival compared to those inoculated with the captive microbiome. Full-length 16S rRNA gene sequencing revealed successful establishment of diverse taxa from *E. spinigera*, including *Dysgonomonas, Wohlfahrtiimonas*, and *Scrofimicrobium*, some showing strong association with either the larval gut or frass. Notably, enrichment with the *E. spinigera* microbiome suppressed a pathogenic *Pseudomonas aeruginosa* strain that dominated the captive microbiome and reduced larval survival in mono-inoculation trials. These findings demonstrate that microbiome enrichment from wild relatives can enhance BSFL health and resilience, supporting a top-down approach to probiotic development for industrial insect rearing.

**Importance:** Black Soldier Fly (*Hermetia illucens*) larvae are increasingly farmed to convert organic waste into valuable products like animal feed and fertilizer. Microbiome composition influences larval health and growth, but captive rearing may restrict or alter functional microbial diversity. Our study shows that enriching the microbiome of captive Black Soldier Flies with microbial diversity of a wild relative species (*Exaireta spinigera*) improves larval growth while suppressing a pathogenic bacterium. This work demonstrates, for the first time, the potential for utilizing the diversity of microorganisms associated with the over 3000 global species of soldier flies as a source of beneficial probiotics supporting industrial Black Soldier Fly rearing.

## Introduction

Microbiome engineering offers a promising avenue to improve animal health and performance through deliberate modification of host-associated microbial communities. This strategy is particularly relevant for captively reared animals that can exhibit disrupted microbiomes relative to wild counterparts with reduced functional diversity or altered community composition (Rosshart et al. 2017; McKenzie et al. 2017; Deutscher et al. 2019; Reese et al. 2021; Dallas & Warne 2023). Introducing beneficial microorganisms or entire communities could restore lost functions and produce novel microbiomes optimized to captive conditions and desired performance traits (Rosshart et al. 2017; Ribeiro et al. 2017; Wallace et al. 2019; Motta et al. 2022).

Black Soldier Fly larvae (hereafter BSFL) have emerged as the “crown jewel” of an expanding global insect industry for organic waste recycling and production of valuable products such as larvae as feed and frass as fertilizer (Tomberlin & Van Huis 2020; Athanassiou et al. 2025). BSFL are fast growing and can feed on a broad variety of organic waste streams including food waste, agricultural by-products and manure (Siddiqui et al. 2022). As in other insects, the BSFL microbiome may contribute to digestion of complex substrates, supplementation of essential nutrients and pathogen resistance (Engel & Moran 2013; De Smet et al. 2018; Luo et al. 2023; Xiang et al. 2024; Auger et al. 2025), although knowledge on the individual effects of members of the BSFL microbiota is still in its infancy (De Smet et al. 2018; Athanassiou et al. 2025; Auger et al. 2025).

Current BSFL rearing practices typically leave microbial colonization to chance, relying on microorganisms in the fermenting diet or rearing environment. Studies of the captive BSFL microbiome have shown high compositional variability between rearing facilities and diets (Gorrens et al. 2022; IJdema et al. 2022), although with some recurring taxa including the bacterial genera *Enterococcus, Klebsiella, Morganella, Providencia, Dysgonomonas* and *Scrofimicrobium* (IJdema et al. 2022). Given the likely importance of the BSFL microbiome, this variability of BSFL microbiomes across facilities and diets could result in challenges for industrial rearing, including inconsistent growth rates, unpredictable waste conversion efficiency, and vulnerability to opportunistic pathogens.

Introducing beneficial microbial diversity to captively reared BSFL could address these challenges and enhance the efficiency and consistency of BSFL to convert diverse waste streams (Gorrens et al. 2023; Auger et al. 2025). Gorrens et al. (2023) reviewed different origins for potential probiotics that have been explored in BSFL including from BSFL guts, materials associated with BSFL rearing like frass, established commercial probiotics in use for non-BSF applications, and those isolated from exogenous environmental sources (e.g., soil). The latter introduction of microorganisms from exogeneous environmental sources could introduce novel diversity and functional capabilities into the BSFL microbiome. However, to engineer improved BSFL microbiomes, introduced microorganisms must overcome critical ecological barriers for successful establishment (Albright et al. 2022; Henry & Bergelson 2025): Firstly, different environmental conditions in the target environment give rise to environmental filtering (termed “host filtering” in the case of host associated microbiomes, Mazel et al. 2018), excluding microorganisms maladapted to the novel environment they are introduced to. Secondly, antagonistic biotic interactions with existing microbial communities in the target environment may prevent colonization, for example via antimicrobial metabolites or competition for nutrients from the resident community.

We propose that to overcome these ecological barriers for microbiome engineering in BSFL, the over 3000 global species of other soldier flies (Stratiomyidae; Woodley 2001) represent a so far unexplored but promising exogenous microbial resource. Their diversity of habitats in soil, under bark or in water, and variety of diets including decomposing plants and animal matter, plant roots, and animal dung (Woodley 2001) may result in considerable microbiome diversity, including strains that are currently absent in BSFL populations but could confer novel beneficial functions as probiotics. Phylogenetically closely related species often share similar microbiomes, a phenomenon referred to as “phylosymbiosis” (Brooks et al. 2016). Close phylogenetic relationship of hosts can be linked to similar physicochemical conditions and immune system traits in the gut, selecting for colonization with similar microbiota from environmental species pools (host filtering, Mazel et al. 2018; Moran et al. 2019). This could potentially make wild relatives of BSF a rich source for compatible probiotics able to overcome host filtering. In addition, if soldier fly species share similar resident microbiome composition, individual members of the microbiota transplanted from other soldier fly species into BSFL may be better able to overcome biotic barriers to establishment, as they would encounter similar ecological communities in BSFL compared to their source environment. Transplants of microbiomes across related species with functional benefits have been successfully demonstrated in other animals: Between frog species to increase host thermal tolerance (Dallas et al. 2024), from the bison to the cow rumen increasing nitrogen digestibility (Ribeiro et al. 2017) and from warthogs to pigs increasing Africa Swine Flu Virus resistance (Zhang et al. 2020). In host species with specific and potentially co-evolved microbial associates, transplanted strains may fail to establish in related hosts (e.g., in corbiculate bees, Kwong et al. 2014, 2017) and transplanting communities across related hosts can have negative fitness consequences relative to the endogenous microbiome (Adair et al. 2020; Parker et al. 2021). However, the diversity of microbial communities observed for BSFL on different diets and across different rearing operations (Gorrens et al. 2022; IJdema et al. 2022) suggests a relatively adaptable nature of the BSFL-microbiome association that would facilitate introduction of novel diversity from related host species.

Here, for the first time, we test whether the BSFL microbiome can be enriched with microorganisms from another soldier fly species, or whether host-microbiome interactions in soldier flies are too specific for this to be feasible or have beneficial effects on BSFL survival and growth. As a starting point, we follow a “top-down” microbiome design strategy (Henry & Bergelson 2025), introducing a complex microbial community from a related host species to assess phenotypic effects relative to an endogenous or diet-derived environmental microbiome. The soldier fly *Exaireta spinigera* (Wiedemann, 1830) (Stratiomyidae: Beridinae) (Fig. 2D) was used for a source microbiome to transplant into BSFL. *E. spinigera* is native to Australia with larvae that develop terrestrially in rotting plant matter, similarly to BSFL (Woodley 1995; Nartshuk et al. 2021; Mackillop et al. 2022). While there is currently no data on the *E. spinigera* microbiome or digestive physiology, their diet of diverse decaying plant materials (Nartshuk et al. 2021) suggests they may harbour microbial communities capable of degrading complex plant cell wall polymers including cellulose and hemicelluloses. As this capacity is of high interest for industrial BSFL rearing (De Smet et al. 2018; Xiang et al. 2024), we selected *E. spinigera* as a promising microbiome transplant source with the potential to confer benefits to BSFL development.

To experimentally test the colonization success and effects on larval phenotypes of different source microbiomes, we exposed axenic neonate BSFL larvae to different microbiome transplants. Specifically, we compare the established microbiome and phenotypic effects from inoculation with (1) an endogenous captive BSF microbiome from a line bred for over 3 years in captivity, (2) an exogenous (*E. spinigera*) derived soldier fly microbiome, and (3) a mixed inoculation competing the two. We include (4) an inoculation with a non-insect associated environmental microbiome of fermenting organic material (here: chicken feed, used as experimental diet) simulating unspecific inoculation from only dietary sources as may be experienced by industrially reared BSFL and (5) a germ-free (axenic) control as baselines to compare larval phenotypes against.

We use this approach to test the following hypotheses:

1. Transplanted microbiota from *E. spinigera* can establish in BSFL either alone or in competition with endogenous BSFL microbiota.
2. Enriching the microbiome of captive BSFL with microbiota from *E. spinigera* larvae improves larval survival and development.

Using PacBio HiFi full length 16S rRNA gene sequencing, we furthermore establish which bacterial taxa (if any) from *E. spinigera* can form close associations with the BSFL larval gut or frass.

## Methods

### BSF rearing and production of axenic larvae

Black Soldier Flies were reared at 27°C, 70% rel. humidity with a larval diet of chicken feed (Watson & William Budget Layer Pellet Chicken, Australia) with 70% water content. The captively bred line originated from larvae collected out of a compost bin in suburban Brisbane (QLD, Australia) in August 2020 (see Bose et al. 2023) and had been maintained continuously in captivity for 3.5 years at the time of the experiment. Adult flies were kept in a netted cage (47.5 cm x 47.5 cm x 93 cm) under a 14 h: 10 h light: dark cycle with artificial light (SPR AGTECH Black Soldier Fly Breeding LED 50W; Evo Conversion Systems, Texas, USA).

Axenic first instar larvae were produced modifying the protocol in Gold et al. (2020): Egg rafts were collected one day after egg laying. After transfer to a biological safety cabinet, they were placed at the bottom of a sterile Corning® 100 µm nylon mesh cell strainer. Eggs were then surface sterilized by sequential immersion into (1) 70% ethanol, (2) 0.6% sodium hypochlorite, (3) 70% ethanol, (4) 0.6% sodium hypochlorite for 2 minutes each in the wells of an 8 well tissue culture plate, transferring the cell culture insert between each solution with a sterile forceps. To ensure dispersion of individual eggs in the sterilizing solutions, egg clumps were dislodged in step (1) by repeatedly agitating the solution through aspirating and dispensing with a 1 ml sterile pipette tip. Eggs were then washed twice in sterile phosphate buffered saline (PBS). Sterilized eggs were re-suspended in sterile PBS and transferred via pipetting to a sterile Phytatray™ cage (11.4 cm × 8.6 cm × 6.4 cm; Sigma; Fig. 1D) with a bottom layer of Vegitone Infusion Broth NutriSelect® Plus (No 41960, Millipore) with 1.5% agar and incubated at 27°C, 70% humidity until hatching. The closed Phytatray™ cage by design allows for air exchange while preventing contamination with external microorganisms. Phytatray™ cages with surface sterilized eggs were checked daily for contaminant growth on the agar or larval hatching. Cages with visible microbial growth were excluded from the experiment.

**Figure 1.**
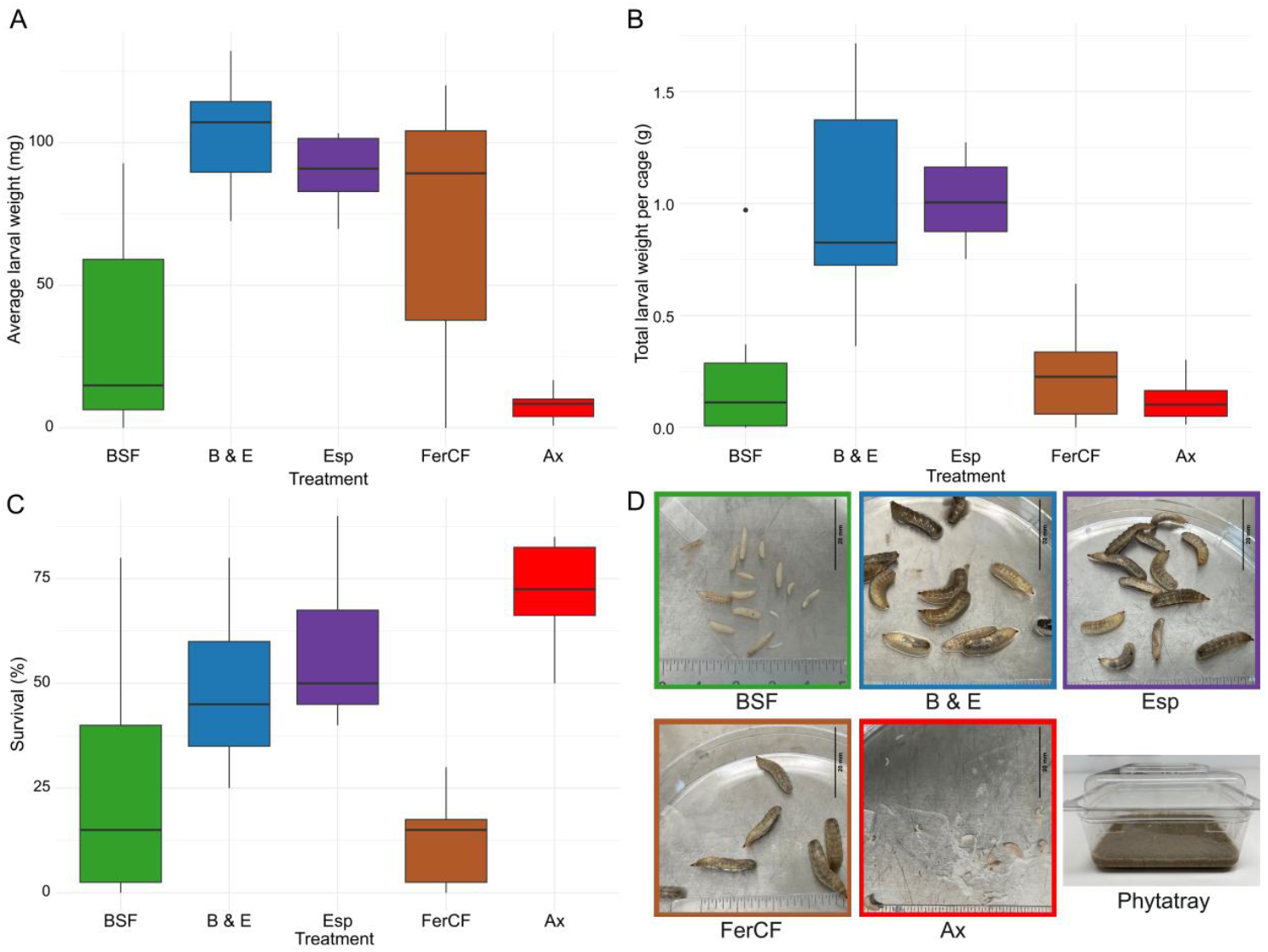
Larval phenotypes across inoculation treatments from transplant experiment. A: Boxplot of average individual larval weight (mg) per container after 2 weeks. B: Boxplot of total larval weight (wet biomass) per container after 2 weeks. C: Boxplot of survival percentages per container. D: Exemplar images of larval phenotypes for experimental treatments (to the same scale) and experimental container (Phytatray™). Inoculation treatments: BSF: captive BSFL microbiome; Esp: Wild *E. spinigera* larval microbiome; B & E: Mixed BSF & Esp microbiome; FerCF: Fermented Chicken Feed; Ax: axenic control.

An axenic experimental diet was prepared by grinding chicken feed (Watson & William Budget Layer Pellet Chicken, Australia) to a fine powder in a conical burr coffee grinder and mixing it at a concentration of 150 g/l with agar (1 g/l) in ultrapure water (Millipore) in aliquots of 500 ml in 1.5 l glass conical flasks. The diet suspension was sterilized via autoclaving at 121°C for 40 minutes. After autoclaving, the diet was cooled down to around 60°C under continuous stirring with a magnetic stirrer. 30 grams of the diet was dispensed into individual Phytatrays™ in a biological safety cabinet and left to solidify for 30 minutes before closing the lid.

### Sourcing and verification of *E. spinigera* larvae

10 *Exaireta spinigera* larvae were collected from a 10-litre plastic bucket with mixed vegetable kitchen waste (12^th^ Oct 2023; CSIRO Black Mountain, ACT, Australia; −35.273442, 149.110524). As the larva of *E. spinigera* has not been described, we ensured selection of *E. spinigera* larvae for the inoculum in the following way: 1.) The timing of the placement of the bucket in spring was chosen to overlap with the activity period of adult *E. spinigera* but precede the summer-active BSF (Mackillop et al. 2022). *E. spinigera* (but no other soldier fly species) was observed visiting and ovipositing in the waste bucket described above. 2.) Larvae from the waste bucket were microscopically examined and found to be morphologically distinct from BSFL due to their shorter, more flattened shape and larger heads with more pronounced, half-spherical eyes protruding from the sides of the head in top view (Fig. 2D; Fig. S1). 3.) The remaining larvae from the bucket after sampling for the inoculum were collected and raised to adults in captivity; only *E. spinigera* adults emerged. 4.) We PCR amplified and Sanger sequenced the cytochrome c oxidase 1 (CO1) barcoding region from extracted DNA of the macerated larval guts (see extraction method below) following the protocol in Folmer et al. (1994) with the primer pair LCO1490 (5’-GGT CAA CAA ATC ATA AAG ATA TTG G-3’) and HCO2198 (5’-TAA ACT TCA GGG TGA CCA AAA AAT CA −3’) and compared the resulting sequence (GenBank accession number PX249732) to published *E. spinigera* sequences.

**Figure 2.**
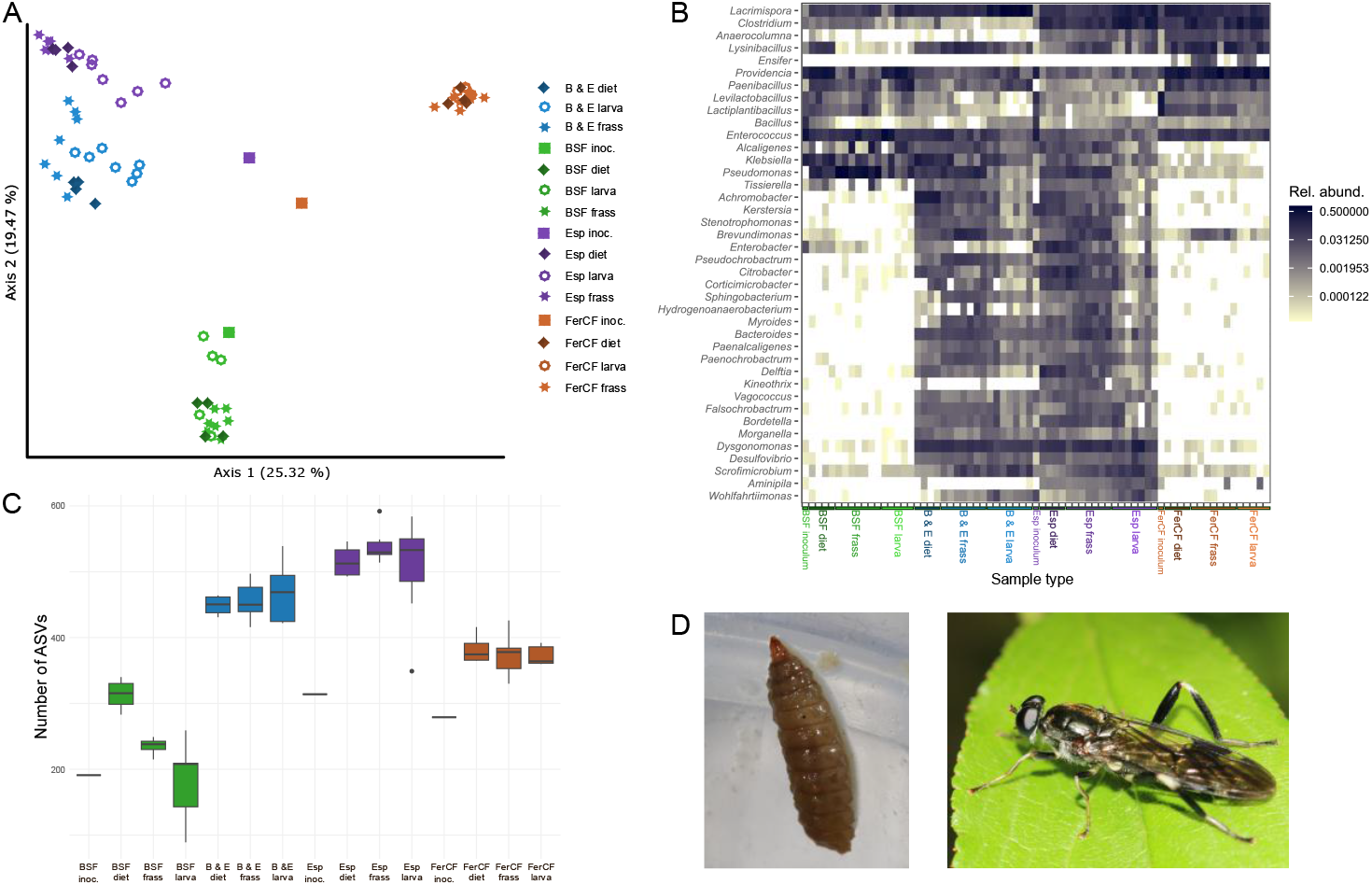
Microbiome composition and diversity of transplant experiment. A: PCoA of Bray Curtis distance matrix for microbial communities (16S rRNA gene ASVs). B: Heatmap of relative abundance (proportion) of top 40 bacterial genera by relative abundance. Heatmap colour scale was log_2_-transformed for better visibility of rare genera. C. Richness (number of ASVs) of microbial communities (rarefied to 25000 reads) across treatments and sample types. D. Larva (left) and adult (right) of *E. spinigera* (images: Hauke Koch). Inoculation treatments: BSF: captive BSFL microbiome; Esp: Wild *E. spinigera* larval microbiome; B & E: Mixed BSF & Esp microbiome; FerCF: Fermented Chicken Feed; Ax: axenic control. Sample types: Larva: dissected larval gut; frass: sample of homogenized frass residue from cages after 2 weeks of larval feeding; diet: sample from inoculated diet after 2 weeks of larva-free incubation; inoc./inoculum: starting inoculum for treatment groups.

### Inoculation treatments

5 different inocula were prepared: 1.) **BSF**: 10 Black Soldier Fly larvae (4^th^ to 5^th^ instar) from the same laboratory colony as above feeding on chicken feed diet (70% water) were dissected to remove whole guts. Guts were combined, weighed, macerated with a plastic pestle in PBS and glycerol (20%) at a 1:10 ratio of gut weight to liquid and stored at −80°C in cryovials. 2.) **Esp:** Whole guts of 10 *E. spinigera* larvae (see above) were dissected out, processed and cryopreserved as in (1). 3.) **B & E**: Equal volumes of the cryopreserved inocula **BSF** and **Esp** were mixed immediately before experimental inoculations of larvae via pipetting combined suspensions up and down 10 times. 4.) **FerCF**: For a baseline comparison of the above treatments with larval phenotypes resulting from inoculation with an environmental, non-soldier fly associated microbiome, 200 mg of 5-day old, fermented chicken feed was macerated and cryopreserved as described in (1). 5.) **Ax**: As axenic control, a sham inoculation with sterile PBS was included. 7 replicate cages were set up for each treatment.

20 neonate larvae (hatched within 24h) were transferred into each of the experimental Phytatray™ cages with sterilized chicken feed agar diets in a biological safety cabinet with a sterilized (0.6% active bleach for 30 minutes, washed in sterile PBS) no 2 sable brush. Inocula 1.) to 4.) were thawed and diluted 1:10 with sterile PBS. 100 µl of the respective 5 inocula were then pipetted over the larvae in each of the cages, immersing the larvae in a fine surface film of the inoculating suspension. Closed Phytatray™ cages were incubated at 27°C, 70% humidity for 2 weeks. Additional control cages of sterile chicken feed diet **without** larvae (4 per inoculum type, excluding the sham inoculum treatment) were inoculated and incubated in the same way as groups (1)-(4) to assess microbiome development in cages in the absence of BSFL.

### Processing of experimental samples

After 2 weeks, all larvae were collected (separately for each cage), surface cleaned by washing in sterile PBS, blotted dry on autoclaved filter paper, and placed into pre-weight 2 ml cryotubes. Larval mass was determined with a microscale to 0.1 mg and larvae snap frozen on dry ice and stored at −80°C. Remaining residual diet/frass mixes in the experimental cages (including in the larva free treatments) were mixed with a sterilized spatula and subsamples of around 1 gram frozen in 2 ml cryotubes at −80°C. For DNA extraction from larval guts, one larva per cage was surface washed with sterile PBS and guts removed via dissection with two pairs of sterile forceps on sterile plastic petri dishes under a dissecting microscope, cleaning the forceps with 80% ethanol followed by heat sterilizing at 250°C with a Steri 250 bead sterilizer (Sigma-Aldrich) and changing petri dishes between each individual larva. For axenic control treatment 5.), surfaces of the agar were washed with 1 ml sterile PBS, and 100 µl of the wash plated out on Vegitone Infusion Broth NutriSelect® Plus (No 41960, Millipore) agar plates (1.5% agar), incubated aerobically for 7 days at 27°C and visually checked for microbial growth. No growth was observed on the plates for 6 of the 7 replicates, but a 7^th^ replicate showed some visible microbial colonies and was excluded from the phenotypic analyses.

### DNA extraction and full length 16S rRNA gene sequencing

DNA was extracted with the DNeasy PowerSoil Pro kit (Qiagen) from the initial inocula and, at the end of the 2-week experiment, one larval gut plus a sample of residual frass from each of the experimental replicates, and samples of the incubated diet from the control cages without larvae. Individual guts or samples of frass / diet (200 mg) were placed into PowerBead Pro tubes of the DNeasy PowerSoil Pro kit together with the provided lysis buffer and a ¼” ceramic sphere bead (MP Biomedicals). Extractions followed the DNeasy PowerSoil Pro kit protocol, with bead beating of sample tubes in a Qiagen Vortex adapter on a Vortex-Genie 2 at maximum speed (= 10) for 10 minutes, and elution after column purification in 100 µl solution C6. We included a control extraction starting with empty lysis buffer and processed in the same manner as the experimental samples.

To characterize bacterial communities, full length 16S rRNA gene sequencing was conducted with HiFi PacBio Revio technology at the Australian Genome Research Facility (AGRF, Brisbane, Australia). The 16S rRNA gene was PCR amplified from extracted genomic DNA using indexed forward F27 (^5’^GCAT/barcode/AGRGTTYGATYMTGGCTCAG^3’^) and reverse primer R1492 (^5’^GCATC/barcode/RGYTACCTTGTTACGACTT^3’^) and sequencing libraries prepared using the Kinnex kit (PacBio). An empty extraction control and one sample of DNA extracted from a frass residue of the axenic chicken feed (**Ax**) treatment were also submitted as control for sequencing, undergoing the same preparation as experimental samples from microbiome inoculated cages. As these latter two samples failed in the 16S rRNA gene PCR amplification step (indicating a lack of intact bacterial DNA), no sequencing libraries could be generated.

### Sequence analysis and taxonomic annotation

Demultiplexed sequence data was processed with the Nextflow PacificBiosciences HiFi-16S-workflow pipeline version 0.9 (https://github.com/PacificBiosciences/HiFi-16S-workflow), filtering for input reads over Q20 and using cutadapt to trim primers (Martin 2011). The maxEE parameter for DADA2 (Callahan et al. 2016) filterAndTrim was set to 2 and pseudo-pooling used for ASV (amplicon sequence variant) detection with a minimum number of samples required to keep any ASV at 1 and minimum number of reads required to keep any ASV at 5.

For diversity analyses, the QIIME 2 (Bolyen et al. 2019) rarefaction curve sampling depth was set to 25000 reads after inspecting alpha rarefaction curves as sequencing depth at which rarefaction curves were saturated and samples could be retained. ASV richness, Shannon entropy and Faith’s Phylogenetic Diversity were calculated for rarefied sample libraries in QIIME2. ASVs were taxonomically annotated in the above pipeline using a naïve Bayesian classifier with the Silva database (Quast et al. 2013; Silva v138.2, Ref NR 99). We also explored taxonomic annotation with GTDB (Parks et al. 2022; GTDB_bac120_arc53_ssu_r220) and GreenGenes2 (McDonald et al. 2024; gg2_2024_09) for comparison. We made the following manual changes to the Silva annotation based on the List of Prokaryotic names with Standing in Nomenclature (LPSN; Parte et al. 2020; accessed July 2025: https://lpsn.dsmz.de/) and our own phylogenetic analyses (see below) of ASVs relative to type strains (see Suppl. Figures S4-S6): A “Koukoulia aurantica” ASV was changed to *Wohlfahrtiimonas* sp. as the former name has not been validly published and the ASV was part of a monophyletic group with described *Wohlfahrtiimonas* species (Fig. S4); all genus “Lachnoclostridium” ASVs were changed to *Lacrimispora* sp. as the former genus is not validly published (Haas & Blanchard 2020) and corresponding ASVs were annotated as *Lacrimispora* by the naïve Bayesian classifier using GTDB or GreenGenes2 as reference (see also Fig. S5); “Actinomyces minihominis” and other closely related *Actinomyces* ASVs were changed to *Scrofimicrobium* sp. following the taxonomic revision in Lao et al. (2025) and our own phylogenetic analyses (Fig. S6).

For phylogenetic analyses, full length 16S rRNA gene sequences of ASVs of interest from this study were combined with sequences of related type strains downloaded from the LPNS database (see above: https://lpsn.dsmz.de/). Sequences were aligned with ClustalW (Thompson et al. 1994) and maximum likelihood phylogenies reconstructed with IQ-TREE 2 (Minh et al. 2020) calculating non-parametric bootstrap support for nodes and using ModelFinder (Kalyaanamoorthy et al. 2017) to choose the best model by AIC (Akaike Information Criterion) score. TreeViewer (Bianchini & Sánchez‐Baracaldo 2024) was used to prepare figures from the IQ-TREE 2 output.

### Microbiome compositional and statistical analyses

Microbial community compositional similarity was visualized with a PCoA using a Bray Curtis distance matrix in QIIME 2 (Bolyen et al. 2019). We used the phyloseq R package (McMurdie & Holmes 2013) to plot the relative abundance (as proportion of total reads) of bacterial genera and selected ASVs across samples via heatmaps. Analyses of Compositions of Microbiomes with Bias Correction 2 (ANCOM-BC2; Lin & Peddada 2020, 2024) were carried out with the corresponding R package to investigate differentially abundant ASVs across sample types. To detect ASVs introduced with the *E. spinigera* transplant that were able to form specific associations with BSFL, we focussed the ANCOM-BC2 on two categories samples from the Esp and B & E treatment groups, conducting a pairwise comparison across communities in larval guts, larval frass and larva-free diet. We pre-filtered the ASVs to those present in at least 35% of samples and performed a Holm method correction of p-values to account for multiple testing. The sample type (gut, frass, diet) and inoculum (Esp, B&E) were incorporated as fixed effects in the ANCOM-BC2 model.

The ANCOM-BC2 result was then filtered for the ASVs most strongly and specifically associated with the larval gut or larval frass. For larval associations, the result of the pairwise ANCOM-BC2 comparison was subset to ASVs that had a significant difference in abundance between larval and frass samples with at least 1 log_e_-fold increase in larvae, and a significant difference between larvae and inoculated diet samples with at least a 2 log_e_-fold increase in larvae. For frass associations, the result of the pairwise ANCOM-BC2 comparison was subset to ASVs that had a significant difference in abundance between frass and larvae samples with at least 2 log_e_-fold increase in frass, and a significant difference between frass and inoculated diet samples with at least a 2 log_e_-fold increase in frass.

### *Pseudomonas aeruginosa* isolation, identification and infection trial

We isolated a strain of *Pseudomonas aeruginosa* (BSFL-1) from a cryopreserved frass sample of the BSF inoculation treatment on Vegitone Infusion Broth NutriSelect® Plus (No 41960, Millipore) with 1.5% agar (grown aerobically at 27°C). Genomic DNA was isolated from an overnight culture with the DNeasy PowerSoil Pro kit (Qiagen), a library prepared with the Oxford Nanopore Rapid Barcoding Kit 96 V14 (SQK-RBK114.96) and sequenced on a PromethION R10.4.1 flow cell (FLO-PRO114M; Oxford Nanopore Technology) at the Biomolecular Resource Facility (BRF) of the Australian National University (ANU). Reads were assembled with Flye v. 2.9.3 (Kolmogorov et al. 2019). The resulting single contig was analysed for the presence of virulence factor genes using the Virulence Factor Database VFanalyzer (Liu et al. 2019), comparing it to *P. aeruginosa* / *P. paraeruginosa* reference strains PAO1, LESB58, PA7 & PA14. The genome and positions of selected virulence factors was visualized in Proksee (Grant et al. 2023) following an annotation with Bakta (Schwengers et al. 2021). One of the 4 identical 16S rRNA gene copies from the genome was aligned with the dominant *P. aeruginosa* ASV found in the BSF treatment and sequences of *Pseudomonas* type strains including *P. aeruginosa* and closely related species from the LPSN database (Parte et al. 2020: https://lpsn.dsmz.de/) for a phylogenetic analysis following methods outlined above. As the recently described *P. paraeruginosa* (Rudra et al. 2022) has an identical 16S rRNA gene sequence to *P. aeruginosa*, we performed a whole genome sequence based taxonomic analysis uploading the genome of *P. aeruginosa* BSFL-1 to the Type (Strain) Genome Server (TYGS), available under https://tygs.dsmz.de (Meier-Kolthoff & Göker 2019; Meier-Kolthoff et al. 2022). Following methods outlined in Meier-Kolthoff et al. (2013; 2022) and implemented by the TYGS, a Genome BLAST Distance Phylogeny (GBDP) was inferred with FastME 2.1.6.1 (Lefort et al. 2015) for our isolate and closely related *Pseudomonas* type strains.

To test the effect of *P. aeruginosa* BSFL-1 on BSFL, we prepared Phytatrays™ with autoclaved chicken feed (150 g/l) and agar (10 g/l) and 20 axenic larvae (replicating methods in the transplant experiment above). 7 replicates each were either inoculated with 100 µl of an *P. aeruginosa* BSFL-1 overnight culture (1:100 PBS dilution of OD 1 culture in Vegitone Infusion Broth (Millipore)) or, in the axenic control, received 100 µl sterile PBS. Phytatrays™ were incubated for 2 weeks at 27°C, 60% rel. humidity, after which live larvae were counted for each replicate. For axenic controls, surface washes of experimental cages at the end of the experiment were plated out on Vegitone Infusion Broth as described above for the transplant experiment. No microbial growth was observed for any of the 7 replicates.

## Data availability

Raw 16S rRNA gene sequencing reads for all samples have been deposited in the NCBI Sequence Read Archive (SRA) under BioProject accession number PRJNA1345878 (SRR35804451-SRR35804521) (to be publicly released). The *Pseudomonas aeruginosa* BSFL-1 genome is available under BioProject PRJNA1313879 (GenBank accession number CP199984). The CO1 barcoding region sequence for the *Exaireta spinigera* larvae serving as microbiome transplant donor is accessible under GenBank accession number PX249732 (https://www.ncbi.nlm.nih.gov/nuccore/PX249732).

## Results

### Phenotypic effects of microbiome inoculation

We observed significant differences in average individual larval weight per cage (ANOVA, F(4, 29) = 10.8, p <.001), total larval weight per cage (ANOVA, F(4, 29) = 14.49, p < 0.001), and larval survival per cage (ANOVA, F(4, 29) = 9.344, p < 0.001) between treatment groups, with the Esp and B & E groups performing consistently better for all three measures compared to the BSF and FerCF inoculated groups (Fig. 1 A-C).

Survival of larvae was highest in axenic larvae on sterilized chicken feed diet (Ax) (average 71.7%) and lowest in larvae receiving the BSF (average 25.7%) and FerCF (average 12.1%) inocula (Fig 1C) with Esp (average 57.8%) and B & E (average 48.6%) inoculated larvae showing intermediate survival. Average individual larval weight was very low for axenic larvae on raw chicken feed (Ax), ranging from ∼1-17 mg (average 7.9 mg) and highest in the Esp (average of 90.3 mg) and B & E (average of 102.8 mg) inoculated larvae. The BSF inoculated group (average of 34.1 mg) had significantly lower average larval weight than the B & E (Tukey’s HSD, p = 0.002) and Esp groups (Tukey’s HSD, p = 0.015) (Fig. 1a). FerCF had a wide range of average individual larval weights, but not significantly different from the Esp (p = 0.74) and B & E (p = 0.30) groups. Total larval mass per container was significantly lower in the BSF group (average 0.24 g) compared to Esp and B & E (both average 1.01 g; Tukey HSD, p < 0.001 for both comparisons). Despite having relatively high average individual larval weights, FerCF larvae had similarly low total larval biomass per cage (average 0.24 g) as the BSF group due to low survival, and the axenic group had the lowest biomass per cage out of all treatments (average 0.12 g) (Fig. 1 A-C). Representative larval phenotypes for all treatments are shown in Fig. 1D.

### Microbiome composition and diversity

The microbiomes in our experiment showed high consistency in community compositions within treatments, and large differences for replicates receiving different inocula (Fig. 2a, b). The microbiomes found developing from the same inoculum on diet in cages devoid of larvae, or in the frass and larval gut were generally compositionally similar. Overall, the mixed inoculation treatment (B & E) was more similar to the Esp inoculated group than to the BSF group (Fig 2a, b).

Plotting the relative abundance of the 40 most dominant genera showed over 20 genera that were either absent or very rare in the BSF and FerCF group but consistently present in the Esp and B & E group (Fig. 2b), including *Dysgonomonas, Bacteroides, Morganella, Scrofimicrobium, Wohlfahrtiimonas, Desulfovibrio, Myroides, Vagococcus, Paenalcaligenes, Sphingomonas* and *Citrobacter*. The genera *Kineothrix* and *Aminipila* were consistently present in the Esp group but virtually absent in the mixed inoculum B & E group. *Clostridium* and *Anaerocolumna* were more common in the FerCF and Esp groups. *Providencia, Enterococcus* and *Lacrimispora* were well represented across all groups, with the BSF group in addition harbouring higher levels of *Pseudomonas*, and the FerCF group *Lysinibacillus*.

This lack of generic diversity in the BSF and FerCF group was also reflected in the lower total number of ASVs per sample compared to the Esp and B & E group (Fig. 2c), as well as lower Shannon entropy and Faith’s phylogenetic diversity (Fig. S2 & S3). Together this showed that BSFL inoculated with a microbiome transplant from our captively bred BSF colony had depauperate microbiomes, but substantial diversity was introduced from the *E. spinigera* larval inoculum, both in isolation and when mixed with the BSF inoculum.

### Strain level associations

The ANCOM-BC2 analysis revealed several ASVs significantly enriched in larval guts compared to frass and the microbiomes developing on inoculated diet in larva-free cages, including two *Dysgonomonas* ASVs, a *Wohlfahrtiimonas* ASV (see Fig. S4), two *Scrofimicrobium* ASVs, a *Vagococcus* ASV, and 20 *Lacrimispora* ASVs (Fig. 3A; S8). The strongest indication for a specific association with growth in the larval gut out of the ASVs above was found for the *Wohlfahrtiimonas* ASV, one of the *Scrofimicrobium* ASVs and the two *Dysgonomonas* ASVs that all had around a 5-6 log_*e*_ −fold reduction (i.e. >99%) in estimated relative abundance on the larva-free diet compared to the larval gut, indicating a very limited ability to persist without the BSFL host (Fig. 3A). These ASVs were absent in the BSF treatment and inoculum (Fig. S8) but present across the Esp treatment and in the Esp inoculum and therefore can be assumed to have originated from *E. spinigera*. The *Scrofimicrobium* ASVs in the experiment belonged to several distinct lineages, with the ASVs strongly associated with the larval gut as distinct lineages branching basally from the phylogeny relative to described *Scrofimicrobium* species, three additional ASVs forming a clade related to *Scrofimicrobium canadense*, and one ASV as sister taxon to *Scrofimicrobium minihominis* (Fig. S6). The 20 *Lacrimispora* ASVs associated with the larval gut belonged to 3 major clades on the 16S rRNA gene phylogeny (Fig. S5): Two ASVs were close to *L. celerecrescens* and *L. sphenoides*, three were close to *L. indolis*, and 15 were close to *L. xylanolytica* and *L. aerotolerans*, indicating substantial diversity of this genus in association with BSFL in the experiment. The ASVs close to *L. indolis* and *L. celerecrescens*/*L. sphenoides* were absent in the BSF group, suggesting an introduction with the Esp inoculum (Fig. S8). In contrast, the ASVs close to *L. xylanolytica*/*L. aerotolerans* were mostly found across all four (BSF, Esp, B&E, FerCF) inoculated treatment groups suggesting a presence in all inocula.

**Figure 3.**
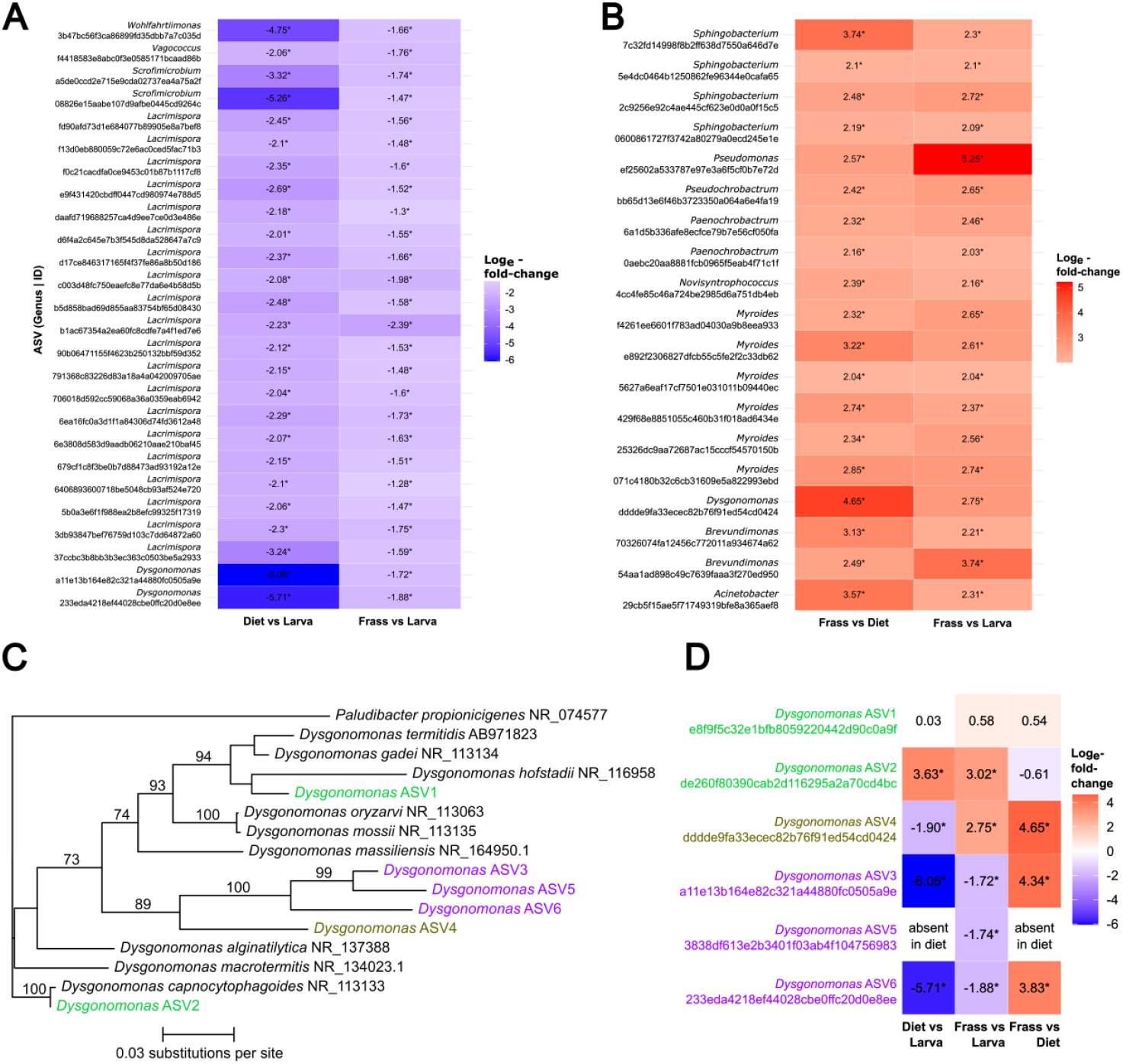
ASV level associations with BSFL. Sample types: Larva: dissected larval gut; frass: sample of homogenized frass residue from cages after 2 weeks of larval feeding; diet: sample from inoculated diet after 2 weeks of larva-free incubation. A: ASVs significantly associated with the larval gut, i.e., ANCOM-BC2 analysis finding significantly higher abundance in the larval gut relative to frass and to microbiome growing on diet in larva-free cages. B: ASVs significantly associated with frass, i.e., significantly higher abundance in the frass over larva free diet, and significantly higher relative abundance in frass over larval gut. C: 16S rRNA gene-based phylogeny of 6 most abundant *Dysgonomonas* ASVs and all *Dysgonomonas* type strains. Bootstrap support on top of branches. Label colours indicate type of association with BSFL (see Fig. 3D). Purple: Specifically associated with larval gut; brown: Specifically associated with larval frass; green: no close association with larval gut or frass. D: Log_e_-fold change in relative abundance (ANCOM-BC2) of 6 top *Dysgonomonas* ASVs from the larval gut relative to larva free diet and frass and from diet to frass.

Several ASVs were overrepresented in the larval frass and significantly underrepresented in the inoculated larva free diet, suggesting that they could specifically grow in larval frass. This group included ASVs from the genera *Dysgonomonas, Acinetobacter, Brevundimonas, Myroides, Paenochrobactrum, Pseudochrobactrum, Pseudomonas* (see Fig. S9), *Sphingobacterium* and *Novisyntrophococcus* (Fig. 3B; S8). Within *Myroides*, 6 different ASVs were associated with frass that formed 3 distinct clades in a phylogeny with related type strains, one close to *Myroides guanonis*, and two closely related to *Myroides fluvii* and *Myroides odoratus* (Fig. S7). *Myroides* was absent in the BSF treatment group and inoculum, but present in the Esp group and inoculum, suggesting an introduction of the above ASVs from *E. spinigera* to the BSFL in the experiment (Fig. 2B; S8).

As the genus *Dysgonomonas* was the most consistently abundant one among the bacterial genera introduced through the *E. spinigera* inoculum (Fig. 2B), we investigated the distribution of the most abundant 6 *Dysgonomonas* ASVs separately. These *Dysgonomonas* ASVs showed remarkably different distributional patterns across larval gut, frass and larva-free diet samples (Fig. 3D): While *Dysgonomonas* ASV1 occurred throughout all 3 sample types with no significant association, *Dysgonomonas* ASV2 was relatively rare in the larval gut, but significantly enriched in the frass and larva-free diet. ASVs 3, 5 & 6 were absent or very rare in larva-free diet and enriched in gut versus frass samples. ASV4 was also rare in larva free diet but enriched in frass versus larval gut samples. The 16S rRNA gene sequences for the BSFL gut or frass associated ASVs 3, 4, 5 & 6 formed a monophyletic clade with around 10% sequence divergence to the closest described *Dysgonomonas* species (Fig. 3C), indicating that this may represent a novel, insect associated *Dysgonomona*s clade.

### *Pseudomonas aeruginosa* BSFL-1 characteristics and infection trial

A single *P. aeruginosa* ASV was notably abundant in all BSF inoculated cages (but rare or absent in the other treatments), with a median relative abundance of 55.5% of reads in frass samples, and 7.1% in larval guts (Fig. 4A; Fig. S10). While this ASV was also present in all 7 replicates of the B & E treatment (showing it had been consistently introduced with the initial inoculum), only very low levels well below 1% rel. abundance were found in frass or larval gut samples of this group (Fig. 4A; Fig. S10). Different *Pseudomonas* ASVs were present in the B & E and Esp groups affiliated with *P. palmensis*/*P. putida* and *P. paralcaligenes* (see phylogeny Fig. S9 and heatmap Fig. S10).

**Figure 4.**
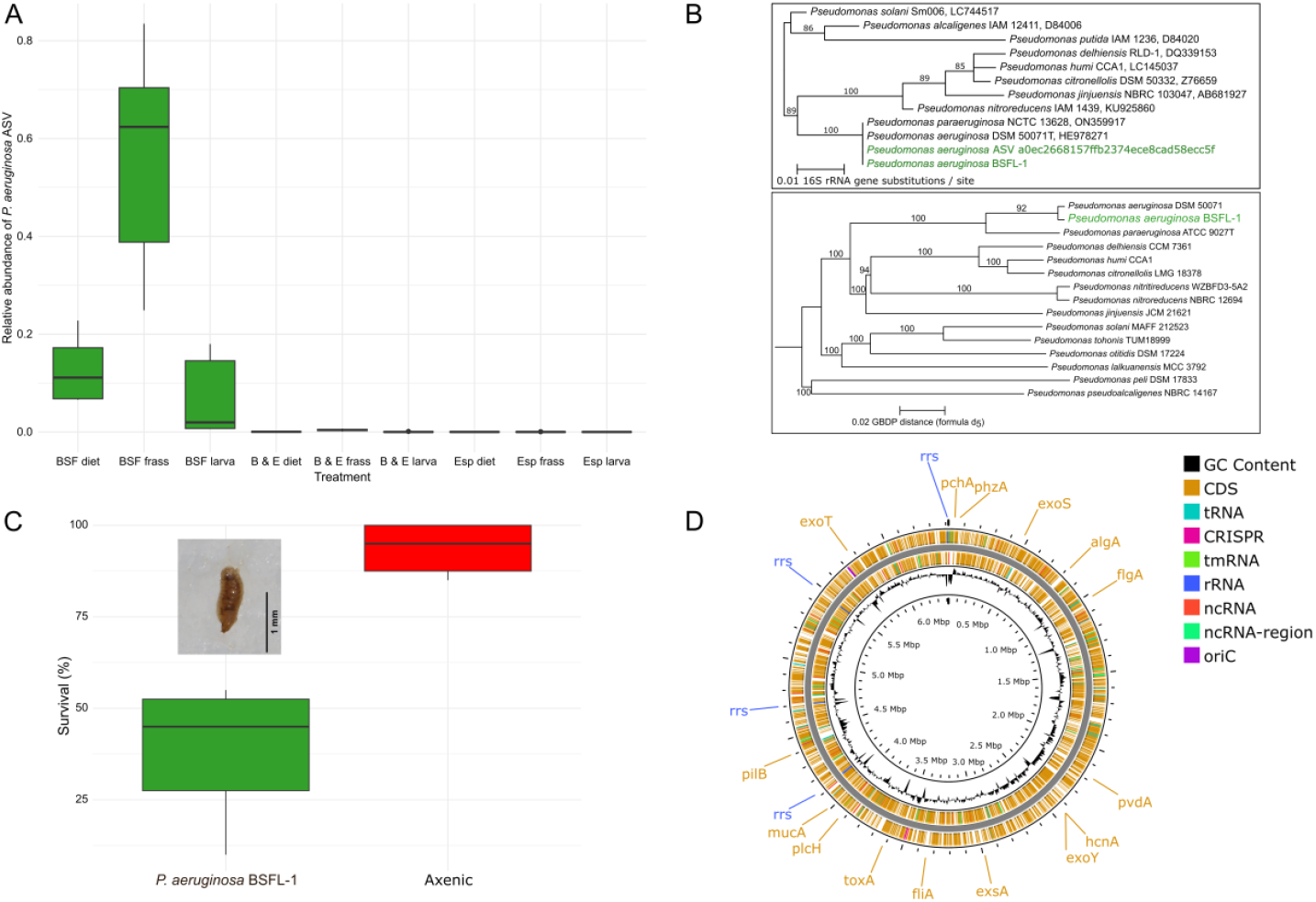
*P. aeruginosa*: A: Boxplot of relative abundance (proportion of reads) of the *P. aeruginosa* ASV dominating the BSF treatment, but rare in the B & E treatment. B: Top: 16S rRNA gene phylogeny (ML) of the *P. aeruginosa* ASV, *P. aeruginosa* BSFL-1 isolate (both in green) and related *Pseudomonas* type strains (strain ID and GenBank accession number after species name). Bootstrap support (100 replicates) on branches. Bottom: Midpoint rooted Genome BLAST Distance Phylogeny (GBDP) of *P. aeruginosa* BSFL-1 (green) and related type strains from the Type Strain Genome Server (TYGS; Meier-Kolthoff & Göker 2019). Tree inferred with FastME 2.1.6.1 from GBDP distances calculated from genome sequences. Branch lengths are scaled in terms of GBDP distance formula d5. Numbers above branches are GBDP pseudo-bootstrap support values > 60 % from 100 replicates. C: Boxplot of survival percentage (live larvae after 2 weeks) for axenic neonates either mono-inoculated with *P. aeruginosa* BSFL-1 or axenic controls. D: Genome of *P. aeruginosa* BSFL-1. Side labels show selected virulence genes (see Suppl. Table 1) and 16S rRNA gene loci (rrs).

We isolated and sequenced the genome of a strain of *P. aeruginosa* from a frass sample of the BSF experimental group (GenBank BioProject PRJNA1313879). The isolated strain *P. aeruginosa* BSFL-1 has a genome of 6.42 Mb with a single, circular chromosome (GC content 66.4%) (Fig. 4D). The genome has 4 identical 16S rRNA gene copies that 100% matched the sequence of the highly abundant *P. aeruginosa* ASV in the BSF frass and larval gut samples (Fig. 4B). A Genome BLAST Distance Phylogeny (GBDP) further identified the strain as “classical*” P. aeruginosa*, and not the close relative *P. paraeruginosa* with identical 16S rRNA gene sequence (Rudra et al. 2022) (Fig. 4B).

*P. aeruginosa* BSFL-1 has the typical range of virulence factor genes found in other strains of *P. aeruginosa* in the VFDB (Liu et al. 2019; see Grace et al. 2022) including complete pathways responsible for iron uptake (pyoverdine, pyochelin), anti-phagocytosis (alginate production and regulation), secretion systems (T3SS with effector proteins exoS, exoT, exoY; T6SS) and toxin production (Exotoxin A, hydrogen cyanide) (241 out of 5975 genes in total, Fig. 4D; Table S1).

Neonate BSFL mono-inoculated with *P. aeruginosa* BSFL-1 and reared gnotobiotically on sterile chicken feed had significantly reduced survival after 2 weeks compared to axenic control larvae (Fig. 4C) (Wilcoxon rank sum test with continuity correction: *W* = 0, p = 0.002; average survival 38.5% for *P. aeruginosa* inoculated vs. 93.6% for axenic). In *P. aeruginosa* BSF-1 inoculated cages, multiple dead first instar larvae showing signs of melanisation were found on the walls of the cages, while no dead larvae with these symptoms were seen in the axenic cages (example image Fig. 4C).

## Discussion

Our findings demonstrate that the microbiome of captive BSFL can be enriched with the microbiome of a wild relative (*E. spinigera*) resulting in improved larval growth and survival. In particular, a strain of the bacterium *P. aeruginosa* shown here to be pathogenic to BSFL and proliferating in the low diversity microbiome from our captive BSFL colony was supressed through the introduction of an *E. spinigera* microbiome transplant. The suppression of opportunistic pathogens by resident host microbiota has been termed “colonization resistance” and documented in a range of hosts, including mammals (Buffie & Pamer 2013) and insects like locusts and bees (Dillon et al. 2005; Koch & Schmid-Hempel 2011). Higher diversity of the resident microbiota can increase colonization resistance (Dillon et al. 2005; Henry & Bergelson 2025). This pattern was also found in our experiment for the B & E group, for which an increase in the depauperate diversity of the microbiome from captive BSFL by the introduction of *E. spinigera* microbes was linked to a near complete suppression of the pathogenic *P. aeruginosa* BSFL-1. *P. aeruginosa* has broad pathogenic potential including in humans and flies (Apidianakis & Rahme 2009). It has previously been demonstrated to induce 80% mortality in 5th instar BSFL larvae upon injection into the hemolymph (Klüber et al., 2022), although BSFL can also express antimicrobial peptides with activity against *P. aeruginosa* (Van Moll et al., 2023). Tegtmeier et al. (2021) previously showed that isolates of *Wohlfahrtiimonas larvae, Morganella morganii, Alcaligenes faecalis* and *Bacillus velezensis* originating from the BSFL gut had activity against *P. aeruginosa* in an *in vitro* zone of inhibition test. The first two genera were also present in the *E. spinigera* inoculated treatment groups (but absent in the BSF inoculated group) in our *in vivo* experiment and could have contributed to the protective effect. However, controlled *in vivo* experiments with isolated strains of the *E. spinigera* derived microbiome diversity will be needed to elucidate the roles of individual introduced bacterial strains to evaluate their potential use as probiotics.

Suppression of pathogenic bacteria may be most important for BSFL at the neonate or first instar stage, which appears most vulnerable with mortality rates of 20-60% (Woods et al. 2019), likely due to an underdeveloped immune system. Consistent with this, we observed high mortality of neonates feeding on *P. aeruginosa* contaminated diet, suggesting that an early intervention via pathogen suppressing probiotics for BSFL neonates in captivity may be key. Bacterial diseases are poorly studied in BSFL but could represent currently overlooked or future problems for the BSF industry (Jensen & Lecocq 2023). For example, in a recent study of industrially reared BSF, infections with the bacterium *Paenibacillus thiaminolyticus* induced high mortality in larvae (She et al. 2023). Furthermore, beyond supporting the health of BSFL, supressing pathogenic bacteria in industrial rearing is important for the safety of BSF factory workers and the suitability of BSFL as animal feed (Moyet et al. 2023; Alagappan et al. 2025).

While the improved survival and development of the BSFL receiving *E. spinigera* microbes in our experiment was likely at least in part due to the suppression of a pathogenic *P. aeruginosa* strain, some of the diversity of introduced bacterial taxa may have had additional benefits that resulted in better larval growth in the B & E and Esp inoculated larvae compared to the BSF and axenic group. For example, the genus *Dysgonomonas* (abundant in group B & E and Esp, but virtually absent in BSF) has a high abundance and diversity of enzymes for the degradation of complex plant polysaccharides (cellulose, hemicelluloses) and likely supports digestion of these compounds in other plant feeding insects such as cockroaches and termites (Vera-Ponce de León et al. 2020; Bridges & Gage 2021) and potentially also BSFL (Li et al. 2023; Xiang et al. 2024). Some *Dysgonomonas* strains can also fix nitrogen (Inoue et al. 2015; Nichio et al. 2025) which could facilitate larval growth on protein poor substrates. Intriguingly, we find strains of *Dysgonomonas* in our experiment that display patterns of relative abundance suggesting they occupy very different environmental niches in relation to BSFL, with some strains strongly associated with the larval gut (and largely unable to grow in the absence of larvae), one strain strongly associated with larval frass (but rare in the larval gut and on larva free diets), and other strains showing no association with larval external or internal environments. Previous studies of the BSFL microbiome reporting *Dysgonomonas* have largely been restricted to short read 16S rRNA gene sequencing (IJdema et al. 2022), and, due to the generally slow evolutionary rate of the 16S rRNA gene and therefore low phylogenetic resolution of these short sequence fragments (Moran et al. 2019) could only treat *Dysgonomonas* in aggregate OTUs precluding insights at the strain level (IJdema et al. 2022). Our results show that there is significant undescribed diversity of *Dysgonomonas* species and strains that can associate with internal or external environments of captive BSFL and fill different environmental niches. From an applied perspective, this ecologically diverse genus may then assist BSFL in both the internal digestion of waste, as well as the external processing of either raw waste material, or excreted frass of BSFL.

Several additional bacterial strains of genera other than *Dysgonomonas* including *Scrofimicrobium, Wohlfahrtiimonas* and *Lacrimispora* were successfully transplanted from *E. spinigera* into BSFL and showed signs of specific colonization of the larval gut in our experiment. The genus *Lacrimispora* was present at high relative abundances in all experimental groups and associated with the larval gut, but increased diversity of strains was linked to the *E. spinigera* transplant. *Lacrimispora* are anaerobic bacteria in the family Lachnospiraceae fermenting complex plant polysaccharides (Kobayashi et al. 2024; Zaplana et al. 2024). They have occasionally been reported in BSFL in association with plant fibre-rich diets (Silvaraju et al. 2024; 2025) and known strains can efficiently degrade xylan (Yang et al. 2025), a major component of the fibre in chicken feed (Choct & Annison 1990). Their presence in larvae feeding on chicken feed in our experiment may thus indicate that the BSFL gut represents a suitable anaerobic niche for *Lacrimispora* to grow on plant fibre indigestible to BSFL themselves. Other strains including from the genera *Myroides, Sphingobacterium* and *Paenochrobactrum* were specifically associated with larval frass.

Notably, many of the genera introduced into BSFL from the *E. spinigera* transplant and forming specific associations with the larval gut or frass have previously been recorded in BSFL. This includes *Dysgonomonas* (Klammsteiner et al. 2020; IJdema et al. 2022), *Scrofimicrobium* (IJdema et al. 2022), *Wohlfahrtiimonas* (Lee et al. 2014; Tegtmeier et al. 2021) and *Myroides* (Raimondi et al. 2020). Wild relative soldier fly species therefore can harbour compatible microbial taxa similar to those found in BSFL, and persisting when co-inoculated with BSFL gut bacteria, confirming our initial hypothesis. This opens the possibility of exploring the diversity of wild soldier fly microbiomes to enhance BSFL performance and restore lost or introduce new functions in industrial rearing. Broader microbiome surveys across soldier fly species are needed to understand the patterns of microbiome diversity and identify promising sources for probiotics.

However, while the overall impact of the *E. spinigera* microbiome transplant was positive, survival in this group was lower than in axenic controls, suggesting that some introduced strains may have pathogenic effects. In addition, the observed pattern of specific colonization of the host gut for some bacterial strains would be compatible with either specific beneficial or pathogenic interactions (Hammer et al. 2019). Therefore, future work should isolate and test individual strains to identify those with probiotic potential.

From an applied perspective, our results support a top-down approach as a starting point for microbiome design (Henry & Bergelson 2025), where complex microbiomes are screened for beneficial effects and colonization potential of target hosts. These initial top-down screens can then be followed up by isolation of strains with probiotic potential and optimization via bottom-up design and evaluation of synthetic communities from isolates (Henry & Bergelson 2025). As has been suggested for plants (Raaijmakers & Kiers 2022), we show that wild relatives of captive animals can represent a valuable resource to “rewild” and enhance the microbiome in this manner.

## Conclusion

Our findings show, for the first time, that microbiome enrichment of the captive BSFL microbiome from a wild relative soldier fly (*E. spinigera*) can improve larval growth and survival. This beneficial effect of “rewilding” the BSFL microbiome appeared to be driven, at least in part, by the suppression of the opportunistic pathogen *Pseudomonas aeruginosa*, consistent with the concept of colonization resistance. Additionally, the introduction of diverse bacterial taxa, including potentially functionally relevant genera for complex carbohydrate degradation such as *Dysgonomonas* and *Lacrimispora*, suggests that microbiome enrichment could confer metabolic benefits beyond pathogen suppression. These results support the feasibility of using wild soldier fly microbiomes as a source of beneficial microbes for industrial BSFL rearing.

## Acknowledgements

HK, AMP and MJM were supported by the CSIRO “Microbiomes for One Systems Health (MOSH)” Future Science Platform (FSP). HK was in addition supported by the CSIRO “Impossible Without You” program. We would like to thank Olivia Young for support with BSF rearing, Bryan Lessard for advice on *E. spinigera* ecology and Carolina Correa-Ospina at the Biomolecular Resource Facility of the Australian National University for bacterial genome sequencing and the Australian Genome Research Facility (AGRF) for carrying our 16S rRNA gene sequencing.

