## Supplemental Figures S1-S10 for "Microbiome enrichment from a wild relative improves Black Soldier Fly larval survival, pathogen suppression and growth"

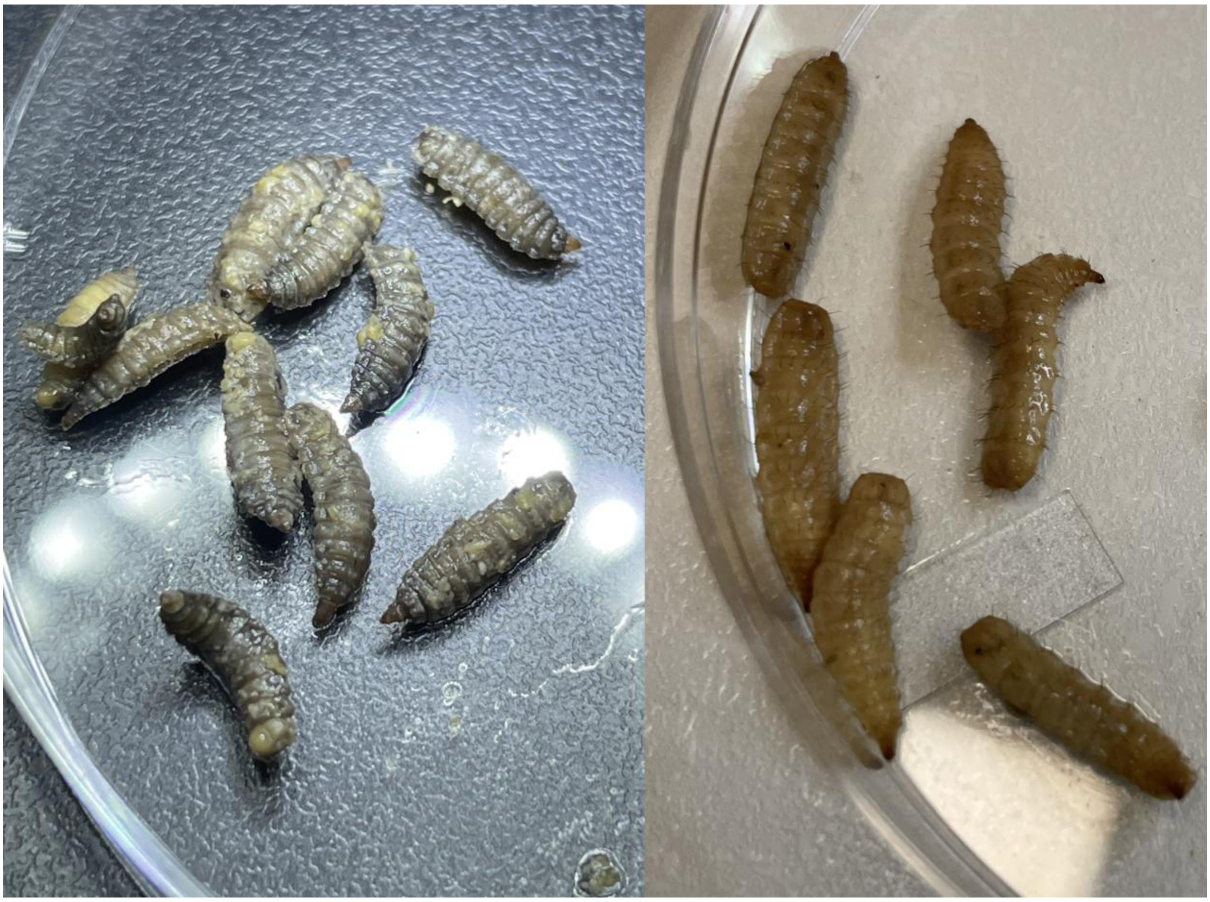

**Figure S1.** Larvae of *Exaireta spinigera* (left) and *Hermetia illucens* (right) used for microbiome transplants.

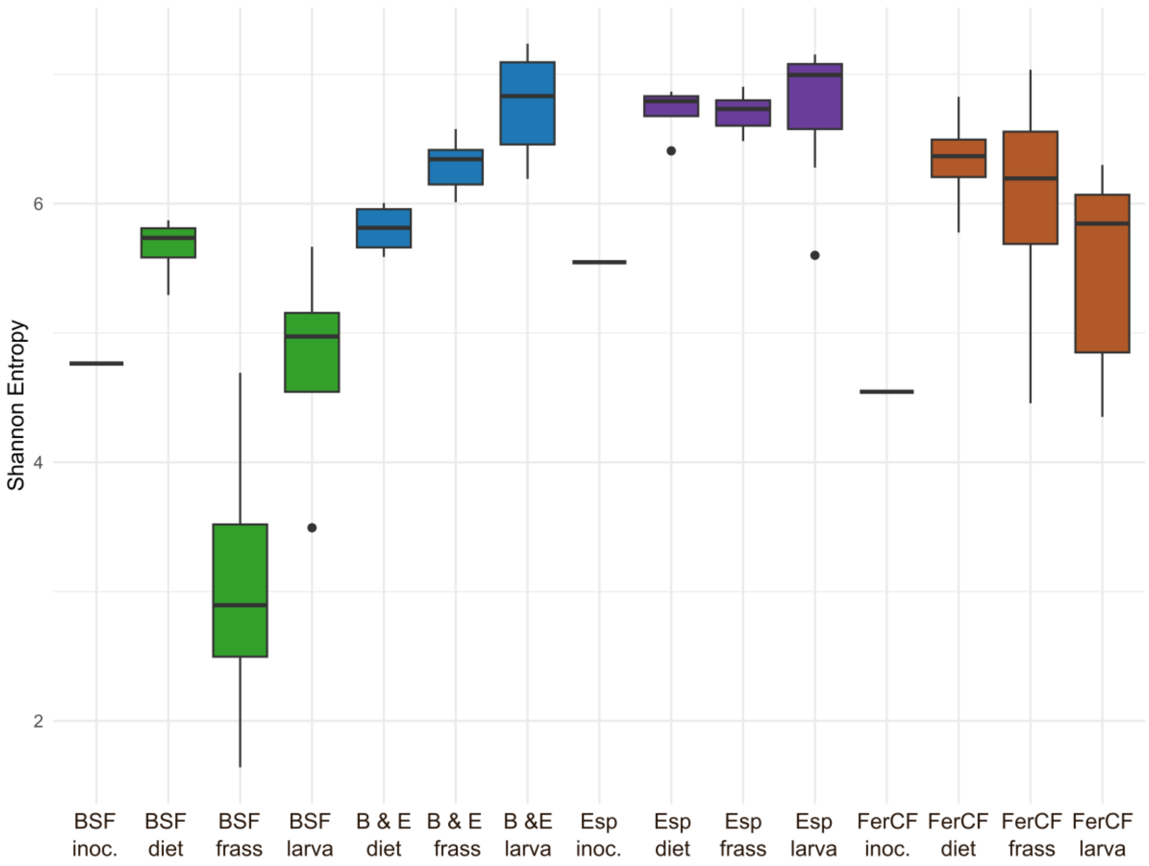


**Figure S2.** Boxplot of Shannon Entropy for ASVs from all sample types, rarefied to 25000 reads per sample.

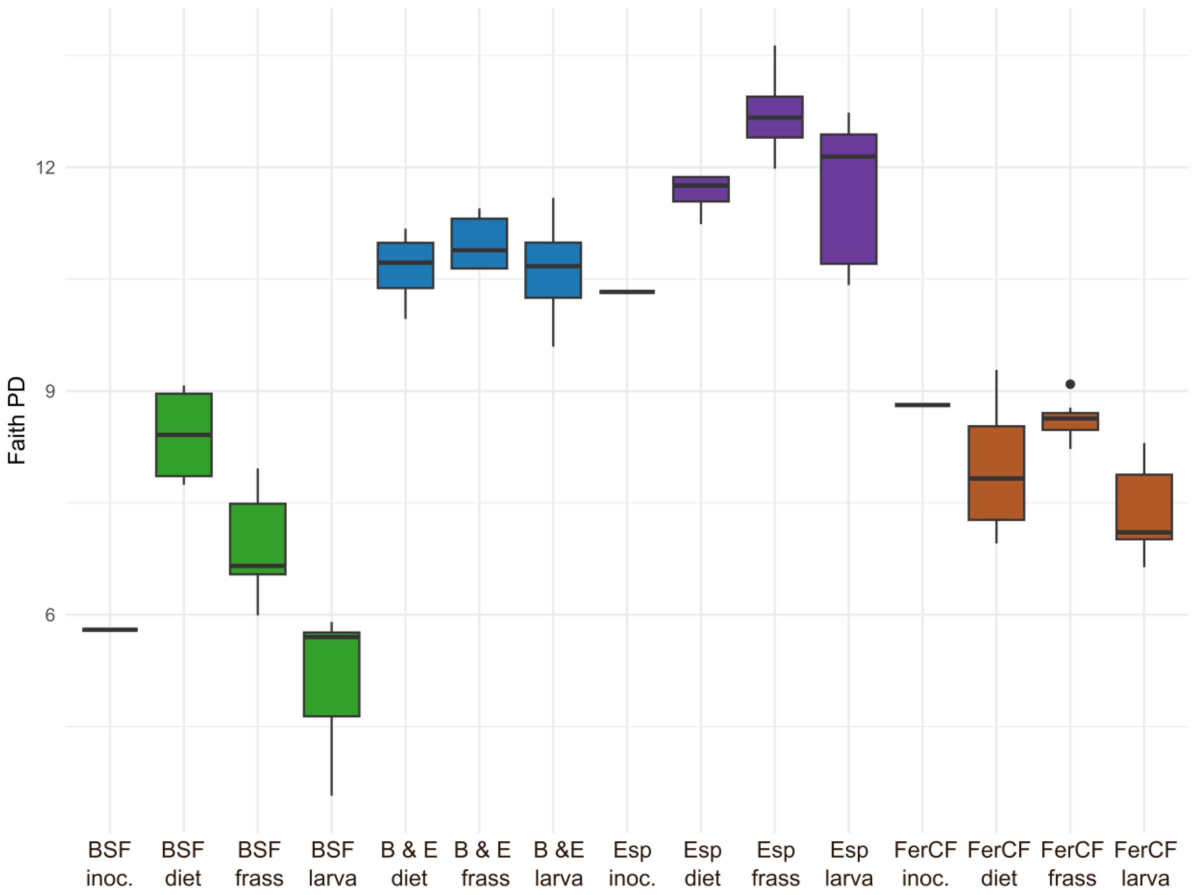


**Figure S3.** Boxplot of Faith's Phylogenetic Diversity for ASVs for all sample types, rarefied to 25000 reads per sample.

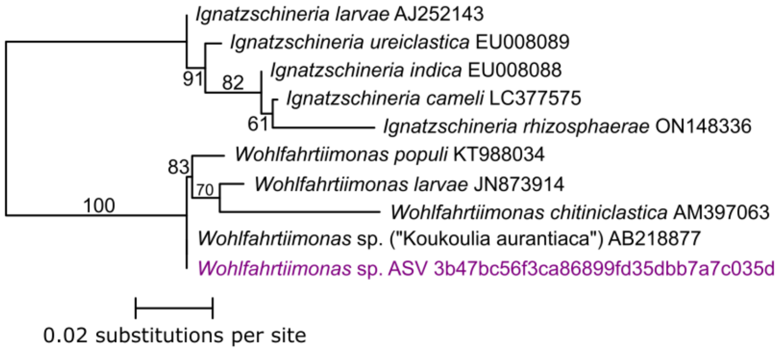


**Figure S4.** Maximum likelihood phylogenetic tree for the 16S rRNA gene of the *Wohlfahrtiimonas* ASV with specific association to the larval gut (purple, see ANCOM-BC2 analysis results) and all *Wohlfahrtiimonas* sp. and closely related *Ignatzschineria* sp. type strains. Bootstrap values over 50 (100 replicates) on branches. ASV identifier or GenBank accession number at the end of tip labels.

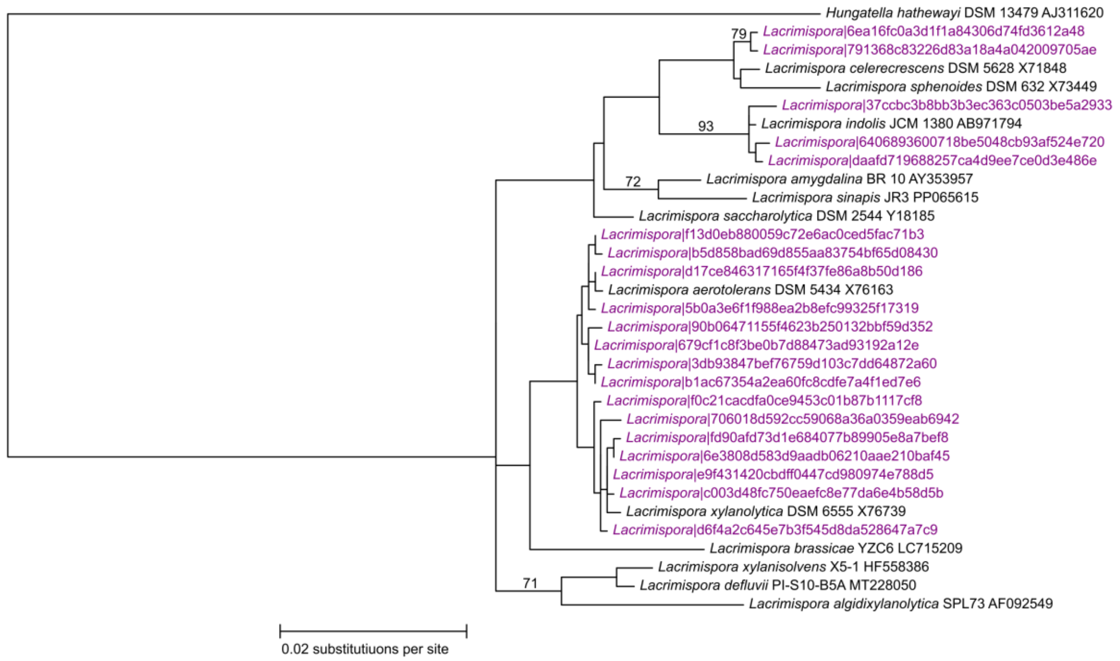


**Figure S5.** Maximum likelihood phylogenetic tree for 16S rRNA genes of *Lacrimispora* sp. ASVs with specific associations to the larval gut (purple, see ANCOM-BC2 analysis results) and all *Lacrimispora* sp. type strains. Bootstrap values over 50 (100 replicates) on branches. ASV identifier or strain name and GenBank accession number at the end of tip labels.

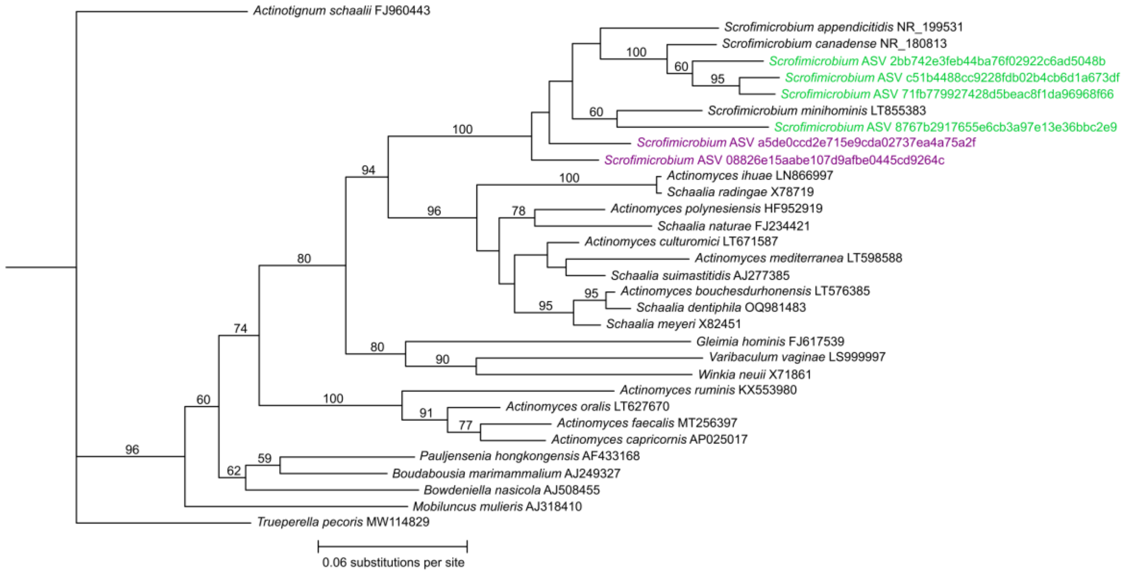


**Figure S6.** Maximum likelihood phylogenetic tree for 16S rRNA genes of *Scrofimicrobium* sp. ASVs with specific associations to the larval gut (purple, see ANCOM-BC2 analysis results), other ASVs (green) and related actinomycete type strains. Bootstrap values over 50 (100 replicates) on branches. ASV identifier or GenBank accession number at the end of tip labels.

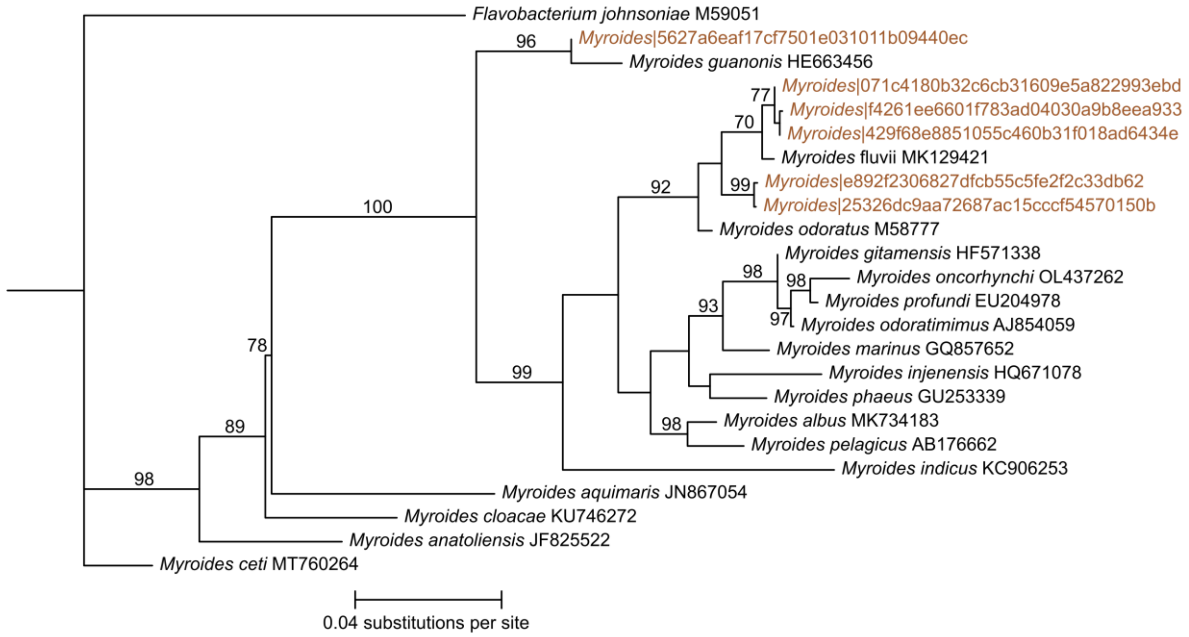


**Figure S7.** Maximum likelihood phylogenetic tree for 16S rRNA genes of *Myroides* sp. ASVs with specific associations to larval frass (brown, see ANCOM-BC2 analysis results) and all *Myroides* sp. type strains. Bootstrap values over 50 (100 replicates) on branches. ASV identifier or GenBank accession number at the end of tip labels.

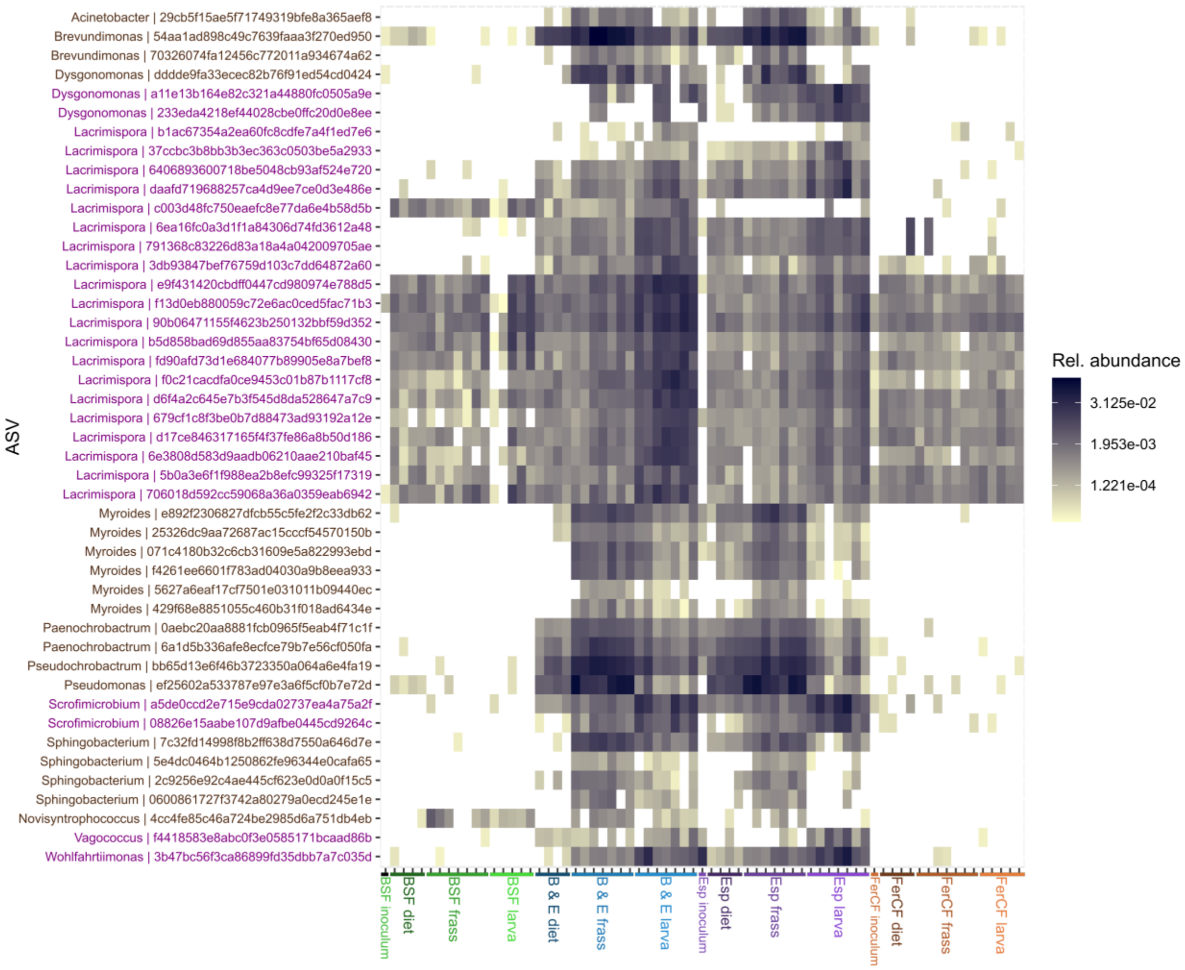


**Figure S8.** Heatmap of relative abundance (proportion of reads) for all ASVs reported in Figure 3 as specifically associated (ANCOM-BC2) with the larval gut or frass across all samples (grouped by sample type). Purple: Associated with larval gut; Brown: Associated with larval frass. $Log_2$ transformed colour scale.

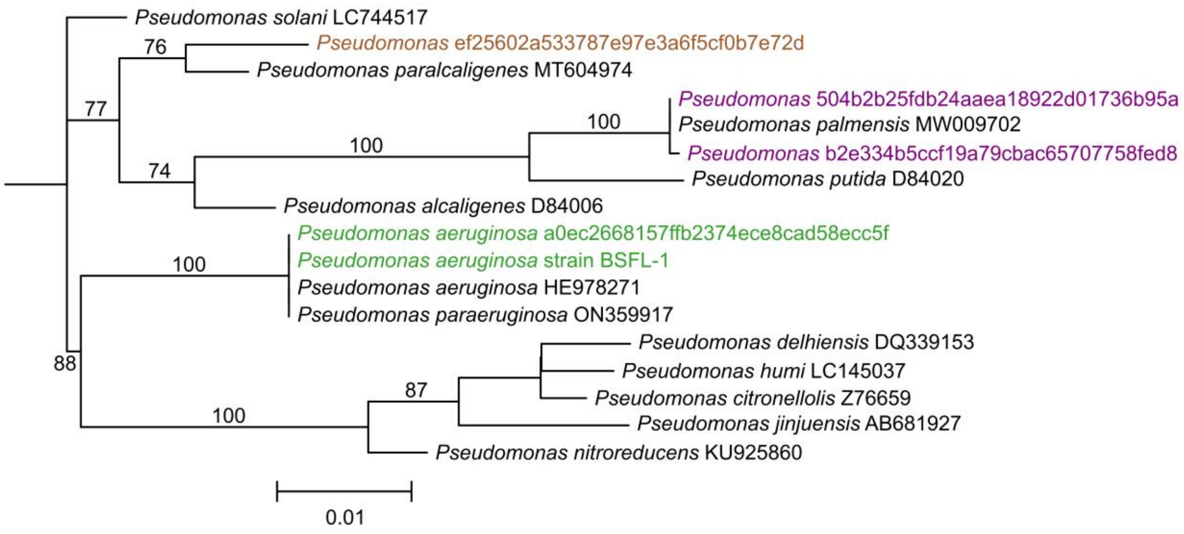


**Figure S9.** Maximum likelihood phylogenetic tree for 16S rRNA genes of *Pseudomonas* sp. ASVs out of top 300 ASVs by relative abundance and including selected, closely related *Pseudomonas* sp. type strains. Bootstrap values over 70 (100 replicates) on branches. ASV identifier or GenBank accession number at the end of tip labels. Green: pathogenic *Pseudomonas aeruginosa* ASV and isolate BSFL-1 associated with BSF treatment; brown: Pseudomonas sp. ASV associated with larval frass (see ANCOM-BC2 analysis Fig. 3); purple: other *Pseudomonas* sp. ASVs mostly associated with Esp and B & E treatment.

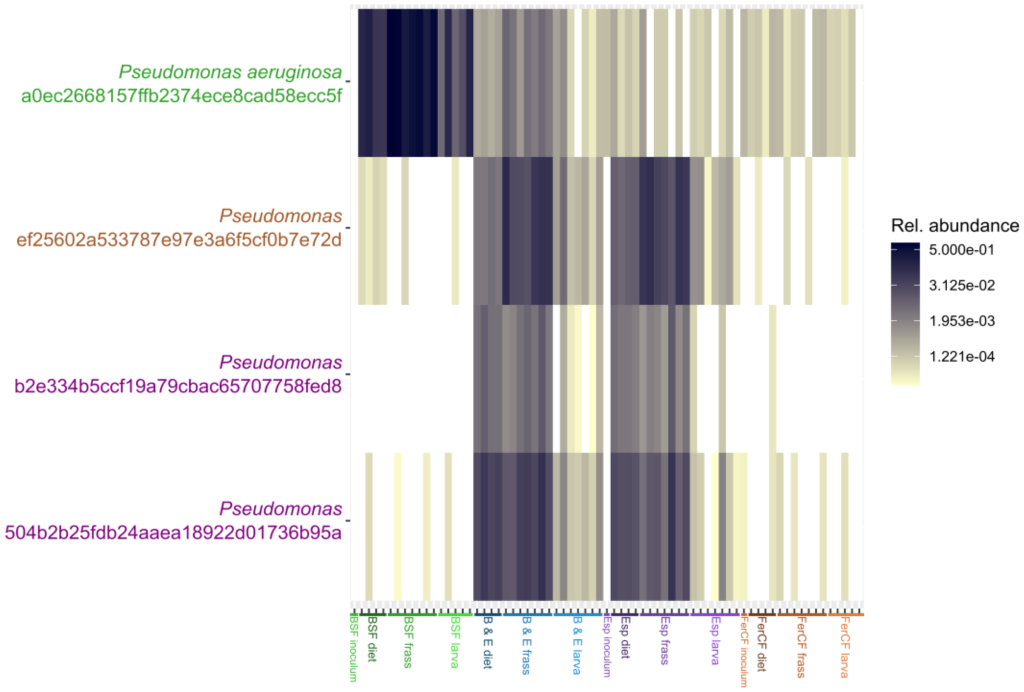


**Figure S10.** Heatmap of relative abundance (proportion of reads) for *Pseudomonas* sp. ASVs from top 300 ASVs by relative abundance across all sample, grouped by sample type. $Log_2$ transformed colour scale. Green: pathogenic *Pseudomonas aeruginosa* ASV associated with BSF treatment; brown: *Pseudomonas* sp. ASV associated with larval frass (see ANCOM-BC2 analysis Fig. 3); purple: other *Pseudomonas* sp. ASVs mostly associated with Esp and B & E treatment; see also Fig. S8.
